# The role of Smarcad1 in retroviral repression in mouse embryonic stem cells

**DOI:** 10.1101/2023.05.18.541392

**Authors:** Igor Bren, Carmit Strauss, Sharon Schlesinger

**Author notes:** Correspondence: Dr. Sharon Schlesinger.

## Abstract

**Background:** Moloney murine leukemia virus (MLV) replication is suppressed in mouse embryonic stem cells (ESCs) by the Trim28-SETDB1 complex. The chromatin remodeler Smarcad1 interacts with Trim28 and was suggested to allow the deposition of the histone variant H3.3. However, the role of Trim28, H3.3, and Smarcad1 in MLV repression in ESCs still needs to be fully understood.

**Results:** In this study, we used MLV to explore the role of Smarcad1 in retroviral silencing in ESCs. We show that Smarcad1 is immediately recruited to the MLV provirus. Based on the repression dynamics of a GFP-reporter MLV, our findings suggest that Smarcad1 plays a critical role in the establishment and maintenance of MLV repression, as well as other Trim28-targeted genomic loci. Furthermore, Smarcad1 is important for stabilizing and strengthening Trim28 binding to the provirus over time, and its presence around the provirus is needed for proper deposition of H3.3 on the provirus. Surprisingly, the combined depletion of Smarcad1 and Trim28 results in enhanced MLV derepression, suggesting that these two proteins may also function independently to maintain repressive chromatin states.

**Conclusions:** Overall, the results of this study provide evidence for the crucial role of Smarcad1 in the silencing of retroviral elements in embryonic stem cells. Further research is needed to fully understand how Smarcad1 and Trim28 cooperate and their implications for gene expression and genomic stability.

**Take home:**

- Depletion of Smarcad1 impairs retroviral repression.
- Smarcad1 is necessary for proper recruitment of Trim28 and H3.3 deposition.
- Depleting Smarcad1 and Trim28 results in enhanced derepression of the MLV provirus.

## Background

The replication of murine leukemia virus (MLV) is restricted in mouse pluripotent cells, namely, embryonic stem cells (ESCs) (1, 2). A complex alliance of factors orchestrates the transcriptional suppression of the viral promoter, known as the long-terminal repeat (LTR), by establishing and perpetuating the repressive chromatin state of the proviral DNA. A pivotal player in this process is Trim28 (tripartite motif-containing 28), also known as Kap1 or Tif1b, which facilitates the recruitment of chromatin modifiers to proviral DNA (3).

Trim28 recruitment is facilitated by ZFP809, a DNA binding protein with zinc finger domains, which recognizes and binds a specific sequence in the provirus called the proline primer binding site (PBSpro)(4). Once Trim28 is associated with proviral DNA, it assembles a coalition of factors involved in transcriptional repression and heterochromatin formation (5). In particular, this ensemble includes SETDB1 (also known as ESET), an H3K9 methyltransferase responsible for methylating histone H3 lysine 9 (H3K9me3) at the proviral DNA, leading to the formation of heterochromatin (6, 7). Consequently, the provirus is packed in a repressive chromatin state, which efficiently blocks the transcriptional machinery from accessing the viral promoter in the LTR. (8). Other components of the silencing complex, such as the YY1 cofactor (9) or the heterochromatin protein HP1 (10–12), contribute to the preservation of the repressive chromatin state and prevent the transcription machinery from reaching the proviral DNA. Similar mechanisms limit the expression of most endogenous retrovirus (ERV) repeats (6, 13, 14). Interestingly, the histone 3 variant H3.3 is also involved in establishing heterochromatin within retroviral sequences. Depletion of H3.3 leads to reduced marking of H3K9me3, suppressing ERVs and adjacent genes (15, 16). H3.3 exhibits a dynamic exchange with the soluble pool of nucleoplasmic histones (17, 18), a phenomenon that is enhanced in ESCs (19–21). H3.3 is important for maintaining ESC pluripotency by regulating gene expression programs central for lineage specification (22–24). Following integration, the MLV provirus exhibits hyperdynamic H3.3 exchange accompanied by transcriptional repression, implying the functional involvement of H3.3 in establishing and maintaining silencing (25, 26).

Another prominent member of the silencing complex is Smarcad1, a nucleosome remodeler that plays a critical role in the regulation of chromatin. Smarcad1, part of the SWI/SNF family of ATP-dependent chromatin remodeling enzymes, is required to maintain genomic integrity and establish repressive chromatin (27). Smarcad1 has been shown to induce nucleosome disassembly and reassembly, suggesting that it plays a role in the dynamic regulation of chromatin structure (28). Smarcad1 also significantly regulates transposable elements (TEs), particularly ERVs, in embryonic stem cells, as its depletion leads to derepression of these elements (29). Moreover, Smarcad1 interacts directly with Trim28 through its CUE1 (CUE domain containing 1) and RBCC protein domains (ring finger and B-box type 2 and coiled-coil domain), respectively (30). Furthermore, Smarcad1 has been suggested to evict nucleosomes, generating accessible DNA crucial for properly recruiting Trim28 and the deposition of H3.3 in Trim28-repressed retroviral sequences (16).

In this study, our objective was to investigate the specific role of Smarcad1 in the silencing of MLV upon ESC infection. Our findings demonstrate that Smarcad1 localizes to the provirus after MLV infection and plays a crucial role in the establishment and maintenance of MLV repression in mouse ESCs. Trim28 recruitment to the provirus requires the presence of Smarcad1, along with H3.3 deposition. Intriguingly, simultaneous depletion of Smarcad1 and Trim28 results in increased derepression of the MLV. These observations suggest a close interconnection between Smarcad1 and the mechanisms used by the silencing complex to suppress proviral transcription, potentially contributing to long-term silencing by dynamically regulating chromatin structure.

## *Materials* and Methods

### Cell lines and cell culture

The cell lines used in this study were KH2 mouse embryonic stem cells, HEK293T and NIH3T3. Cells were passaged every 3-4 days by washing with PBS and adding trypsin EDTA solution. The growth medium for HEK293T and NIH3T3 cells consisted of high glucose Dulbecco’s modified Eagle medium (DMEM, BI, 01-055-1A) supplemented with 10% fetal bovine serum (FBS), 2 mM L-glutamine (BI, 03-020-1A), 100 units/ml penicillin (BI, 03-031-1B), and 100 µg/ml streptomycin (BI, 03-031-1B). ESCs were cultured on 0.2% gelatin-coated tissue culture plates with high glucose DMEM, 15% FBS, 2 mM L-glutamine, 100 units/ml penicillin, 100 µg/ml streptomycin, 200 mM MEM nonessential amino acids (Rhenium, 11140-035), 1 mM sodium pyruvate (Rhenium,11360039), and 0.12 mM β-mercaptoethanol. This medium also contained 2i+LIF: leukemia inhibitory factor (LIF) 1000 units/ml, PD0325091 1 µM (PeproTech, PD 0325901) and CHIR99021 3 µM (PeproTech, 2520691). hygromycin B 140 µg/ml (ENZO, ALX-380-309-G001) was added to maintain this HA-H3.3 cassette in the genome. To induce HA-H3.3 transcription, the KH2 medium was supplemented with doxycycline 10 µg/ml. For antibiotic selection, the media were supplemented with puromycin 2.5 µg/ml or G418 [neomycin] 500 µg/ml (Rhenium, 11811031). The puromycin and neomycin selection processes needed 2 and 5 days to complete, respectively. All cells were cultured in a humidified incubator at 37°C with 5% CO2. The cells were tested for mycoplasma (Hylabs, KI 5034I) every two weeks.

### shRNA design and cloning

***The*** pLKO.1 (Addgene plasmid #13425) lentiviral vector carrying the neomycin resistance gene was used to express shRNA sequences in targeted cells. After lentiviral infection, antibiotic resistance was used to select for the knockdown cells. Cloning of the shRNA hairpin was performed using T4 ligase (NEB, M0202L) at a ratio of 3:1, insert:vector. shRNA sequences were taken from (29). Sanger sequencing of these amplicons confirmed the presence and successful insertion of the shRNAs into the pLKO.1 vectors.

### Production of lentiviruses/retroviruses

HEK293T cells were used to produce lentiviruses and retroviruses. The cells were co-transfected with pMD2. G (Addgene Plasmid #12259) vector for the VSVG envelope and psPAX2 (Addgene Plasmid #12260) for lentiviruses or ECO2 for retrovirus gag-pol genes. After 48 hours, the medium containing viruses was collected, filtered through a 0.45 µm filter and supplemented with 10 mM HEPES buffer (BI, 03-025-1B) and 12.5 µg/ml polybrene (Merck, TR-1003-G). MLV-based GFP-reporter (pNCA-GFP) vectors were used for retroviral transduction assays (as in (31)).

### RNA extraction, cDNA synthesis, and real-time quantitative PCR (qPCR)

Total RNA was extracted from cells using TRI reagent (Sigma, T9424) according to the manufacturer’s instructions. RNA concentration and purity parameters (260/280 & 260/230) were determined by a NanoDrop spectrophotometer. One microgram of RNA was reverse transcribed using the qScript cDNA Synthesis Kit (Quantabio, 95047-100). To control for genomic DNA impurities in the RNA samples, reactions without reverse transcriptase (-RT) were performed simultaneously. Housekeeping genes Ubiquitin C (UBC) and GAPDH were used to calculate the relative expression level of genes of interest by the ΔΔCT method. Each sample was tested in triplicate. All primers have been previously tested and found to agree with standard curve evaluation and are listed in Supplementary Table S1. All qPCRs were performed on a StepOnePlus™ Real-Time PCR system. To examine the expression level of endogenous retroviruses (ERVs), the RNA was treated with TURBO™ DNase (2 U/µL) (Thermo Fisher, AM2238) prior to cDNA synthesis. –RT controls were included in all assays.

### Flow cytometry

ESCs and NIH3T3 cells infected with GFP-reporter MLV retroviruses were analyzed using a CytoFLEX flow cytometer equipped with a 488 nm laser. A minimum of 100,000 cells were examined per sample. The generated data were further analyzed using FlowJo V.10 software.

### Chromatin immunoprecipitation (ChIP)

ChIP was performed using a previously described protocol (32). Briefly, 2 to 3 million cells were cross-linked using 1% formaldehyde (Sigma, F8775) for 10 minutes at room temperature. The crosslinking process was quenched by adding 120 mM glycine solution (Sigma, G8898). The samples were incubated at room temperature for 5 minutes and then centrifuged for 5 minutes (1,500 rpm, 4°C). The supernatants were aspirated and washed in cold PBS supplemented with protease inhibitor 1:25 (PI, Sigma, 11836170001). Pellets were resuspended in 200 µl ChIP lysis buffer supplemented with PI (1:25), incubated on ice for 15 minutes and sonicated using a Qsonica Q800R2 sonicator. After confirming that DNA fragments suitable for immunoprecipitation (200 – 700 bp) were generated, the sonicates were centrifuged for 10 minutes (8,000 rpm, 4°C). Five percent of each sample was collected and stored at 4°C overnight for further use as input. The remaining supernatant was diluted (1:10) in ChIP dilution buffer, and PI (1:25), 20 µl Magna ChIP™ Protein A Magnetic Beads (Sigma, 16-661) and antibodies (1 µg per 1 million-cell chromatin, all antibodies are listed in Supplementary Table S2) were added to tubes, which were then transferred to 4°C for an overnight incubation with shaking at 10 loops/minute. On the next day, the samples were immunoprecipitated and washed with low-salt, high-salt, LiCl, and TE buffers supplemented with PI (1:25). The samples were resuspended in 100 µl ChIP elution buffer. All samples (immunoprecipitated and input) were then transferred to 62°C for 6 hours and shaken at 300 rpm to reverse the cross links. Extraction of the DNA was performed using a QIAquick PCR Purification Kit (QIAGEN).

### Whole-cell extraction and Western blotting (WB)

Ten million cells (per extract) were resuspended in hypotonic lysis buffer composed of ice-cold Tris pH 7.4, EDTA 0.2 mM, DTT 0.5 mM, protease inhibitor (PI, 1:25) and NaVO4 1 mM and mechanically lysed. High salt buffer composed of Tris pH 7.4, EDTA 0.2 mM, DTT 0.5 mM, and NaCl 1 M was added to the samples, and they were centrifuged for 30 minutes (13,000 g, 4°C). The supernatants containing the extracts were collected and diluted 1:1 in SDS sample buffer. The samples were denatured at 95°C for 10 minutes and loaded on Bolt™ 4 to 12% Bis-Tris, 1.0 mm Mini Protein Gels (Thermo Fisher, NW04120BOX). 20X Bolt™ MES SDS Running Buffer (Thermo Fisher, B0002) was used for high-resolution separation of proteins smaller than 110 kDa, while 20X Bolt™ MOPS SDS Running Buffer (Thermo Fisher, B0001) was used for high-resolution separation of proteins larger than 110 kDa. After separation, the proteins were transferred to a nitrocellulose membrane at 20 V in the presence of transfer buffer composed of Bolt™ Transfer Buffer (20X) 50 ml/L (Thermo Fisher, BT0006), 100 ml/L methanol and water. The membrane was blocked for 25 minutes using 5% skim milk in TBST. Next, the membrane was incubated in the presence of primary antibodies (listed in Supplementary Table S2) and 1% skim milk in TBST at 4°C overnight. The next day, the membrane was washed three times with TBST for 15 minutes per wash. Then, the cells were incubated in the presence of horseradish peroxidase (HRP)-conjugated secondary antibodies for one hour at room temperature. The membrane was washed three times under the same conditions. Detection was performed by Pierce™ ECL Western Blotting Substrate (Thermo Fisher).

### Statistical analysis

Statistical analysis was performed using GraphPad Prism 9.5.0 software. Data are presented as the mean values ± SEMs. Statistical significance was determined using Student’s t test. Statistical significance was considered at p < 0.05. Significance levels are ∗ p < 0.05; ∗∗ p < 0.01; ∗∗∗ p < 0.001.

## Results

### Smarcad1 is localized to the provirus after MLV infection

To examine the hypothesis that Smarcad1 plays a role in MLV repression, we transduced ESCs with an MLV-like vector carrying a GFP reporter (MLV-GFP) controlled by the LTR (as in (8)). ChIP[qPCR performed two days after infection (2 dai) showed eminent enrichment of Smarcad1 in some MLV regions: PBS, TES, and coding regions (Figure 1). Smarcad1 occupancy was not observed in the 5’UTR of the MLV LTR (the 40 nt region), near the negative control region (NCR) of the provirus, suggesting that Smarcad1 is located around the PBS together with other members of the Trim28-dependent retroviral silencing complex. Importantly, Smarcad1 enrichment levels in MLV sequences were higher than those observed for ERV and other Trim28 genomic targets that were previously shown to be bound by Smarcad1 (29, 30).

**Figure 1.**
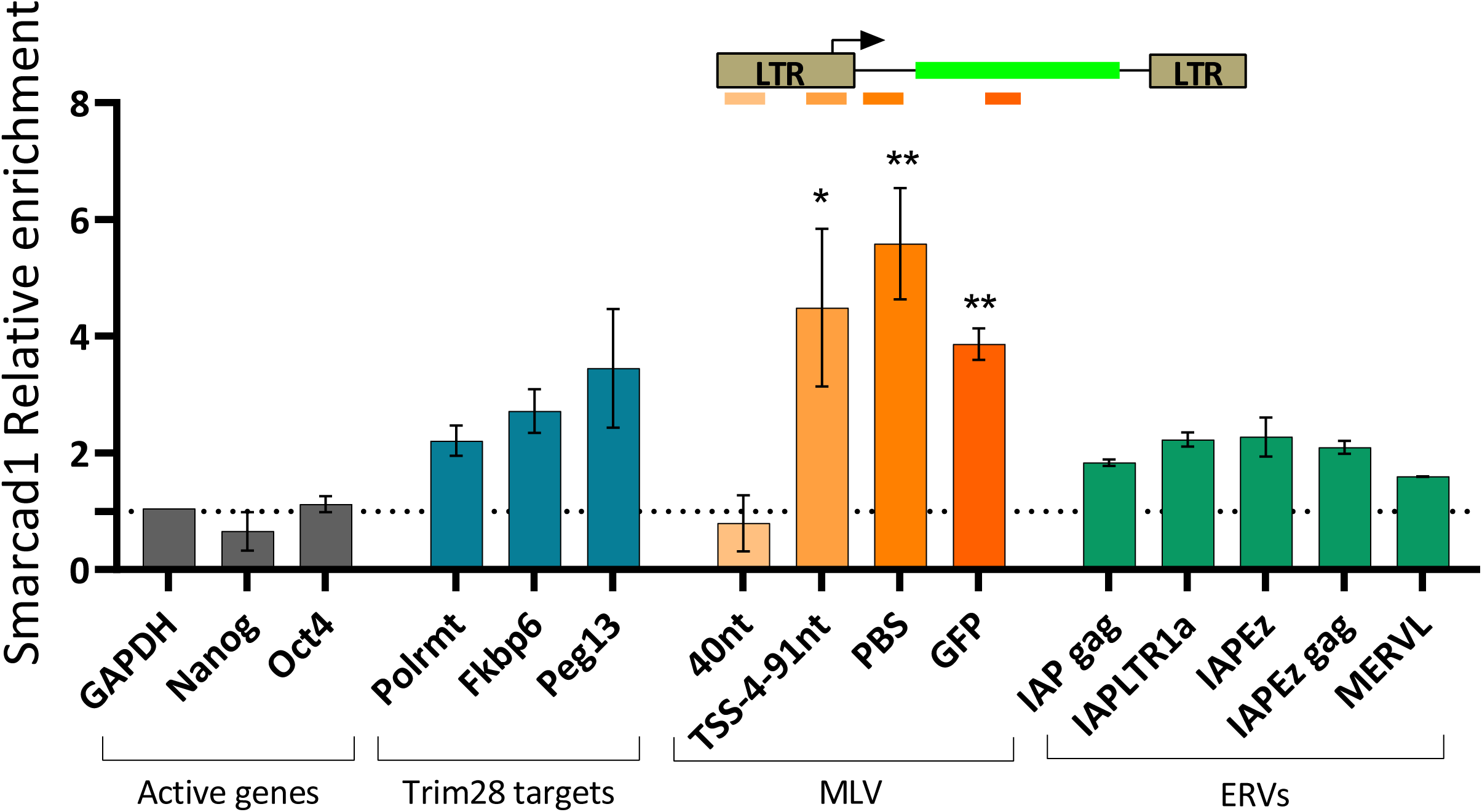
Smarcad1 is recruited to newly integrated retroviruses. ChIP□qPCR was performed using Smarcad1 antibody, 2 days after MLV infection. Primers for open and repressed chromatin loci were used, values shown are % input normalized to nonenriched sites (Gapdh and Oct4). The illustration above the graph indicates the position of the primers on the MLV vector: 40 nt is located by the NCR, at the 5’ end of the LTR, next are the PBS [primer-binding site], TIS [translation initiation site] ang GFP [Green fluorescence protein] primers. n=2, Data are the mean ± s.e.m, asterisk shows significant difference from the negative control genes, P value was calculated using two-tailed unpaired Student’s t test, *P<0.05, **P<0.01.

### Smarcad1 plays a role in the establishment and maintenance stages of MLV repression in mouse embryonic stem cells

To examine whether Smarcad1 enrichment around retroviral TSS has an effect on the transcriptional regulation of the provirus, we depleted Smarcad1 using lentivirus-mediated delivery of short-hairpin RNA (shRNA sequences taken from (29) and cloned and inserted into pLKO.1). KD efficiency was assessed using RT[qPCR (Figure 2A) and verified using Western blotting (Figure 2B and Supplementary Figure S1A). For further experiments, we used shSmarcad1_1 and shControl after we validated the stability of shRNA depletion and normal expression levels of pluripotency-related genes and other key factors related to retroviral epigenetic silencing (Figure 2A and supplementary Figure S1B). Next, we infected the two lines with MLV-GFP and followed the population fluorescence by flow cytometry two and seven days after infection (Figure 2C). Interestingly, no change in GFP expression was found in the less firmly repressed MLV virus, which carries PBSgln and is therefore not as bound by Trim28 (Figure 2D). The observation that the less firm restriction of the MLV vector carrying alternative PBS, namely, PBSgln (10-fold higher GFP expression relative to PBSpro), is not mediated by Smarcad1 (Figure 2D), which is in line with the central role of Trim28 in PBSpro-specific silencing (3, 8) and with the data showing that only PBSpro requires H3.3 for fully efficient silencing (32). Smarcad1 deletion results in a 2.4-fold increase in expression immediately after proviral integration (2dai), which is maintained for several weeks (Figure 2E). Infection and integration efficiency were not affected by Smarcad1 depletion, as there was no change in the number of integrated genomic proviral copies (Supplementary Figure S1C). Therefore, these data suggest that Smarcad1 plays a role in the onset of MLV repression, probably through its binding to Trim28. To examine whether Smarcad1 is also required to maintain MLV repression in ESCs, we first infected KH2 with MLV-GFP and then depleted Smarcad1. A stable elevated GFP signal was observed after Smarcad1 depletion, demonstrating that it is also needed for the maintenance stage of MLV repression (Figure 2F).

**Figure 2.**
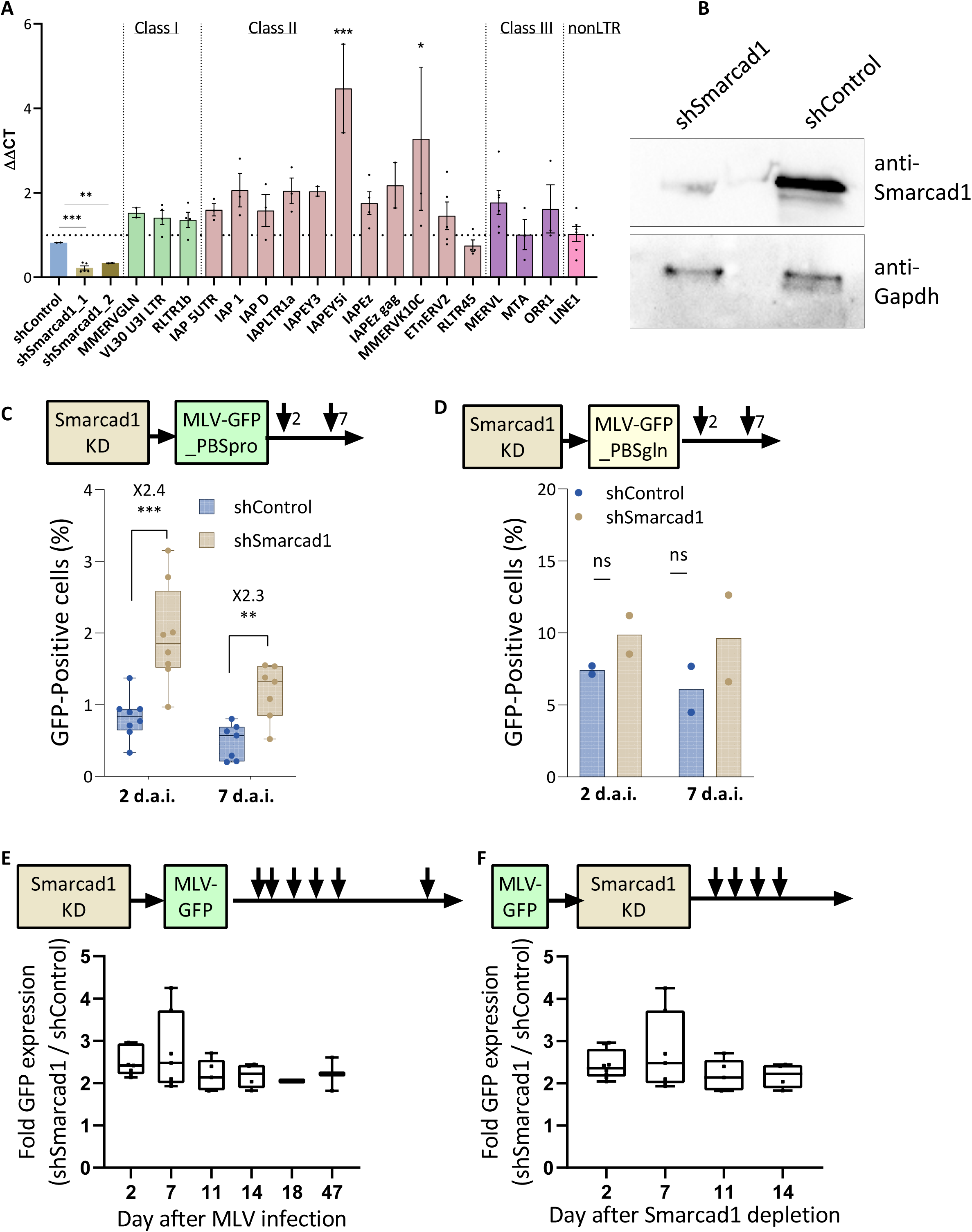
Smarcad1 plays a role in MLV repression. (A) Smarcad1 and ERVs expression changes (shSmarcad1 vs shControl) were measured by RT□qPCR, normalized to UBC control gene. n=2-5, data are the mean ± s.e.m (B) Immunoblotting of shSmarcad1 and shControl ESC extract using Smarcad1 antibody, with anti-Gapdh as loading controls (C) The percentage of GFP-positive cells 2 and 7 days after infection by PBSpro or (D) PBSgln virus in the WT and Smarcad1 depleted ESCs., n=2-6, data are the mean ± s.e.m. (E) Change over time of the GFP-positive cells in the depleted cells vs. the control cells (F) Change over time of the GFP-positive cells in the depleted cells vs. the control cells after infection by PBSpro. n=2-7, data are the mean ± s.e.m. In all panels, P value was calculated using two-tailed unpaired Student’s t test, *P<0.05, **P<0.01, ***P<0.001.

### Smarcad1 is needed for proper Trim28 recruitment and H3.3 deposition in nucleosomes that wrap the MLV provirus after infection

Smarcad1 is an ATP-dependent chromatin remodeling enzyme shown to regulate DNA accessibility in Trim28-binding ERV, possibly by nucleosome eviction (16). Therefore, we hypothesized that in the absence of Smarcad1, the protein complex needed for MLV repression cannot be properly recruited or assembled in the provirus. To further decipher how Smarcad1 depletion disrupts retroviral repression, we first applied ChIP using a Trim28 antibody to WT and Smarcad1 KD cells 2 and 7 days after infection. Trim28 enrichment in the PBS region of the MLV provirus immediately after infection was found to be independent of Smarcad1 (Figure 3A). However, seven days after infection, Trim28 enrichment increased significantly in WT ESCs but not in Smarcad1 KD cells (Figure 3A). Therefore, Smarcad1 is not needed for the initial recruitment of Trim28 to the provirus, but it is important to stabilize and strengthen its binding properly over time. Genomic Trim28 targets, such as the promoter region of Polrmt and Fkbp6 and the ERVK subfamily IAPEz, also lost Trim28 enrichment after Smarcad1 depletion (Figure 3B). Next, since the removal of Smarcad1 was shown to decrease the accumulation of histone variant H3.3 from IAPEz sequences (16), we used KH2 ESCs (33) expressing a single copy of H3.3a-HA controlled by doxycycline (Dox) (34–36) for our H3.3-HA ChIP assays. Exploring the dynamic accumulation of H3.3-HA in the proviral genome 48 hours after infection and 8 hours after Dox induction allowed us to focus on the onset of retroviral silencing immediately after integration. Surprisingly, no H3.3 enrichment was observed in the provirus (Figure 3C), although Trim28 was already bound there at that time (Figure 3A). The accumulation of H3.3 is only observed several days after the integration of MLV and depends on Smarcad1 (Figure 3C). A similar effect of H3.3 eviction following Smarcad1 depletion was observed in Trim28 target sequences such as Polrmt, imprinting control regions such as Peg13, and the IAPEz subfamily (Figure 3D). These data show that although Trim28 recruitment to the PBS is not dependent on Smarcad1, Smarcad1 is important to reinforce the attachment of Trim28 to the area and is essential for H3.3 accumulation in retroviral sequences. These findings are also true for most ERV subfamilies examined here, in which Trim28 binding is barely affected by Smarcad1 depletion, while H3.3 enrichment and heterochromatinization are significantly reduced. Interestingly, the effect of Smarcad1 depletion on ERV class II expression was mild (Figure 2A), and no such effect was observed on MLV-PBSgln proviral expression (Figure 2D). Consistent with this and previously published data, no significant enrichment of Trim28 (Figure 3E) and H3.3 (Figure 3F) was observed in this proviral sequence, and no effect of Smarcad1 deletion was shown. This could be due to other prominent repression mechanisms applied to those sequences (31, 37).

**Figure 3.**
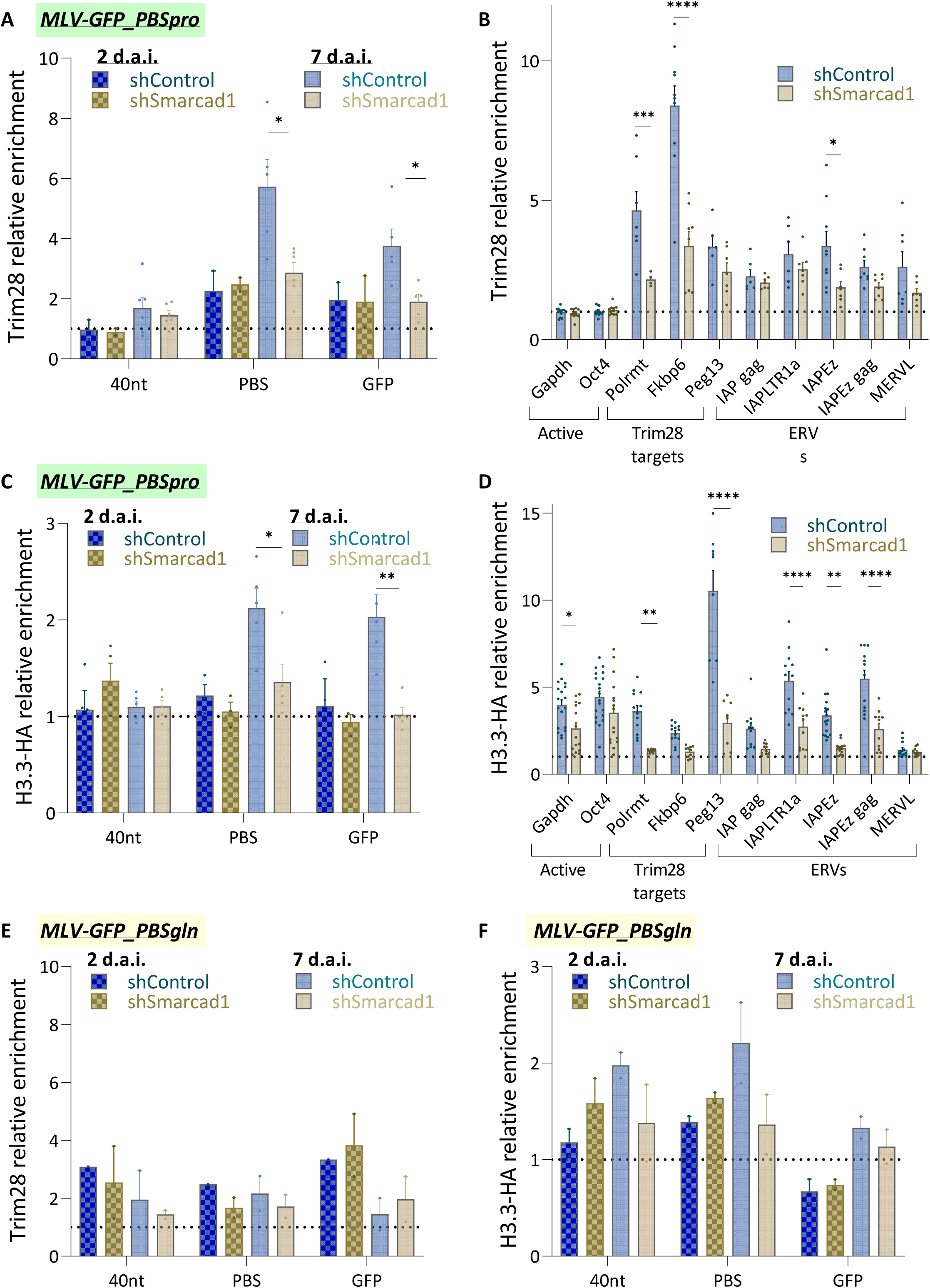
Smarcad1 reinforces Trim28 binding and allows H3.3 accumulation and heterochromatinization of proviral sequences. (A) ChIP□qPCR was performed using Trim28 antibody on day 2 (n=2) and 7 (n=6) after MLV-PBSpro infection, with primers for MLV provirus-specific regions and (B) for open and repressed genomic chromatin loci, including ERVs (n=12). Values shown are % input normalized to nonenriched sites (Gapdh and Oct4). (C) ChIP□qPCR was performed using HA antibody, on day 2 (n=3) and 7 (n=5) after MLV-PBSpro infection, with primers for MLV provirus-specific regions and (D) for open and repressed genomic chromatin loci, including ERVs (n=12). (E) ChIP□qPCR was performed after MLV-PBSgln infection using Trim28 antibody and (F) HA antibody at days 2 and 7 after infection (n=2). Values shown are % input normalized to nonenriched sites. All data are the mean ± s.e.m, in all panels P value was calculated using two-tailed unpaired Student’s t test, *P<0.05, **P<0.01, ***P<0.001, ****P<0.0001.

### The combined depletion of Smarcad1 and Trim28 results in enhanced MLV derepression

Up to this point, we have established that Smarcad1 is required to repress MLV proviruses, selected class II ERVs, and other Trim28-bound genomic loci. Mechanically, we show that Smarcad1 improves recruitment of Trim28 to these sites, allows proper deposition of H3.3 into the nucleosomes that wrap DNA in these genomic regions, and promotes their further marking with H3K9me3 repressive histone marks. To further explore the mechanism leading to MLV repression, we targeted Trim28 from the newly MLV-infected (2d.a.i.) Smarcad1 KD cells using Trim28 shRNA as in (9) and four days of G418 selection and verified double KD by RT[qPCR on day seven after MLV infection (Figure 4A). Double KD did not affect the expression of other chromatin modifiers or key pluripotency genes (Supplementary Figure S2A). Next, we analyzed the cells by flow cytometry for MLV-GFP expression. As expected, Smarcad1 KD and Trim28 KD resulted in ∼2-fold and ∼4-fold increases in the GFP signal relative to shControl, respectively (Figure 4B). Interestingly, the combined depletion of both Smarcad1 and Trim28 resulted in an ∼7-fold increase in the GFP signal relative to shControl, demonstrating an additive effect. These observations were consistent in two independent biological replicates, suggesting that the combined depletion of both regulators results in enhanced derepression of MLV. In addition, the change in the expression level of selected ERVs was examined following the depletion of Smarcad1, Trim28 and the double KD. The upregulation of some ERVs was comparable. Between shTrim28 and double KD cells, others appeared to be more responsive to Smarcad1 depletion (Figure 4C). However, no additive effect of combined depletion is observed for ERV repression. Therefore, we hypothesize a discrepancy in the role of Smarcad1 in the silencing of endogenous and exogenous retroviral sequences. To further test this hypothesis, we repeated the double KD assay in a different order: first, we depleted Smarcad1; second, we infected cells with MLV and kept them in culture for 16 days; and third, we depleted Trim28 and finally analyzed the cells using flow cytometry and qPCR after four days of selection. As before, we measured an ∼2-fold and ∼5-fold increase in GFP signal after depletions of Smarcad1 and Trim28, respectively (Figure 4D), except that this time, the GFP signal observed in double KD cells was comparable to that of shTrim28. RNA data confirmed the depletion of Smarcad1 and Trim28 (Supplementary Figure S2B). These observations suggest that the additive effect of combined Smarcad1 and Trim28 depletion decreases with time and implies that their combined role is more significant in the establishment of MLV repression.

**Figure 4.**
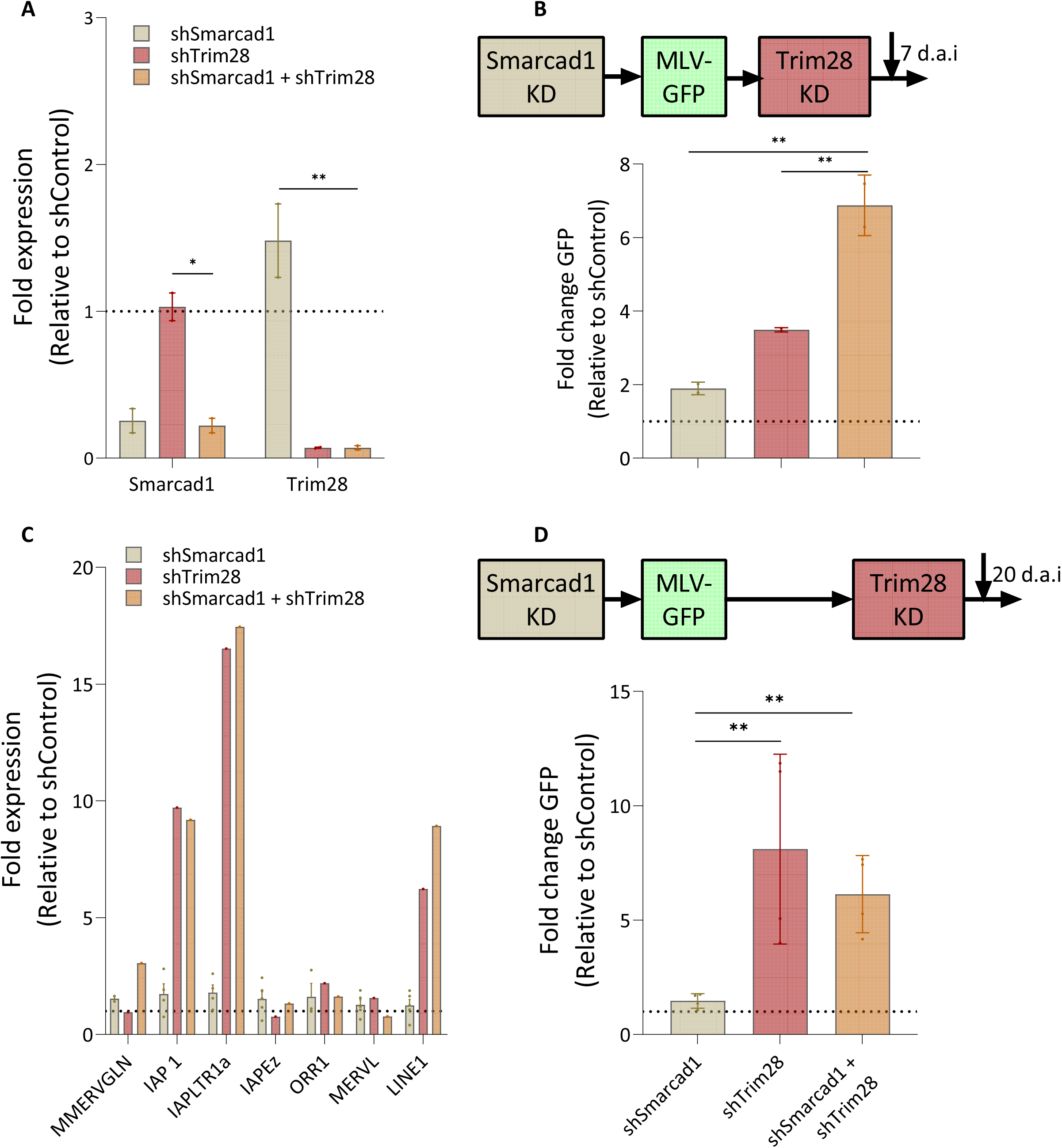
Smarcad1 and Trim28 depletion has an additive effect on MLV upregulation. (A) Double KD was achieved by infection with a lentiviral vector with Smarcad1 shRNA, puromycin selection and then Trim28 shRNA lentiviral vector with G418 selection. Depletion was verified by RT□qPCR, normalized to UBC control gene. Data are the mean ± s.e.m, n=2 (B) The percentage of GFP-positive cells 14 days after shSmarcad1 infection, 7 days after infection by MLV-PBSpro, and 5 days after shTrim28 infection. Data are the mean ± s.e.m, n=2 (C) Expression levels of selected ERVs following Smarcad1 and/or Trim28 depletion were measured by RT□qPCR. Data are the mean ± s.e.m, n=2-5 (D) The percentage of GFP-positive cells 27 days after shSmarcad1 infection, 20 days after infection by MLV-PBSpro and 5 days after shTrim28 infection. Data are the mean ± s.e.m, n=2. In all panels, the P value was calculated using a two-tailed unpaired Student’s t test, *P<0.05, **P<0.01.

## Discussion

We studied the role of the Smarcad1 chromatin remodeler in retroviral repression in the mouse ECS. We introduced a novel temporal dimension by monitoring changes in transcription and epigenetics over time after integrating new proviral sequences into the genome. We found that Smarcad1 played a crucial role in retroviral suppression, both in the early stages and in long-term establishment. Additionally, a correlation was observed between Smarcad1 binding and H3.3 deposition on the provirus and other silenced genomic sites and ERVs. The study also suggests that Smarcad1 had a role beyond supporting Trim28 binding and contributed to silencing independently, particularly in the early stages. Overall, the research revealed the involvement of the Smarcad1-Trim28-H3.3 pathway in repressing newly integrated proviral sequences. Thus, we have acquired fresh perspectives on the mechanisms governing retroviral suppression mediated by Smarcad1. From the point of MLV transduction to cells, in approximately 24 hours, the newly incoming retroviral sequence is being integrated into the cell genome (38). Trim28 is recruited to the MLV provirus via direct interaction with ZFP809 (4), YY1 (9) and EBP1 (39).

Smarcad1 is an important component of the double-strand break repair machinery and is recruited to newly synthesized DNA (27), which could explain its affinity for the newly integrated provirus. Additionally, Smarcad1 and Trim28 proteins also interact directly through their CUE1 and RBCC domains, respectively (30), and Smarcad1 binding is enriched in ERV sequences (29, 30). However, it was not clear whether Smarcad1 also has a role in recruiting Trim28 to newly integrated retroviral sequences. A mutation in the CUE1 domain that abolishes the interaction of Smarcad1 with Trim28 did not impair Trim28 recruitment to its target sites, while the occupancy of Smarcad1 at these loci was reduced (29, 30). Nevertheless, Smarcad1 KD or a mutation in its ATPase domain altered Trim28 recruitment, suggesting that Smarcad1 catalytic activity is needed for proper or enhanced Trim28 binding to its target loci. This is in agreement with our results, which show that while Trim28’s immediate localization to the provirus is Smarcad1 independent, Smarcad1 facilitates the long-term establishment of proviral repression for MLV, as it does for ERV (29). We hypothesize that Smarcad1 plays a role in the reinforcement and stabilization of Trim28 binding to the MLV provirus, as well as to genomic sites. However, two days after MLV transduction, Smarcad1 is essential for complete silencing, suggesting that Smarcad1 could have a Trim28-independent role in the establishment of retroviral repression. Consequently, when both proteins were depleted, the expression levels of MLV-GFP in the double KD cells were higher than in each KD separately, indicating an additive effect of these factors. Therefore, the role of Smarcad1 in silencing, at least in the early stages, is not only to support Trim28-mediated silencing. This is not true for ERV and proviral silencing maintenance, indicating that Smarcad1 plays a Trim28-independent role only at the silencing establishment stage. We speculate that this could be explained by recent findings showing that Smarcad1 associates with key architectural regulatory factors related to genome organization in mammalian nuclei (40).

Is Smarcad1 needed for the accumulation of H3.3 on retroviral sequences? The correlation between Smarcad1 binding and H3.3 deposition on the provirus implies that it is. However, while several studies suggest that H3.3 may influence the local chromatin environment by recruiting chromatin remodeling complexes, particularly SWI/SNF and NuRD (35, 41), a mechanism for Smarcad1-induced H3.3 deposition is less clear. Our data suggest that once Smarcad1 is depleted, H3.3 is demolished. This is in agreement with the idea suggested by (16) that Smarcad1 nucleosome remodeling action is needed for H3.3 deposition. This is also an interesting exception to the general genomic role of Smarcad1 in suppressing histone turnover (42). Recently, Trim28 and H3.3 were shown to interact and regulate MLV silencing (32). While Trim28 is necessary for full H3.3 accumulation on the provirus, H3.3 depletion led to lower levels of Trim28 binding. Here, we show that although retroviral silencing is already prominent on day two after infection (Figure 2C) and (8), no enrichment of H3.3 has yet been observed, suggesting a role for H3.3 in maintaining silencing. Since we also show that H3.3 deposition is Smarcad1 dependent, a mechanism can be proposed. First, Smarcad1 and Trim28 are recruited to the newly integrated provirus, each contributing to some degree to silencing establishment. Next, Smarcad1 allows H3.3 to accumulate on surrounding chromatin, consolidates Trim28 binding, and mediates silencing complex assembly.

Finally, we show that upregulation after Smarcad1 depletion was also stably maintained when Smarcad1 was depleted after MLV integration (Figure 2F), as is the case for endogenous retroviruses (Figure 2A and (29)). Consistently, our analysis of the structure of the chromatin in proviral DNA indicates that similar epigenetic silencing mechanisms are applied to incoming viruses and some ERVs, as previously suggested (8).

Although one limitation of the study is the low enrichment levels of all factors, especially at the two d.a.i time points, the positive control of verified target genes confirms the reliability of all the ChIP data. We also repeated the observation of (29) that different categories of epigenetically repressed loci, including representatives of class II ERVs, imprinted genes, and developmental genes, are all regulated via both Smarcad1 and Trim28 in ESCs. Additionally, following Smarcad1 depletion, enrichment levels of H3.3 were reduced in the same genomic Trim28 target loci, which were highly enriched in control cells. These data imply that a wide array of genomic elements are repressed by Smarcad1-Trim28-H3.3 in ESCs.

In conclusion, this study provides evidence that Smarcad1 is a critical component in the repression of MLV in mouse embryonic stem cells. Smarcad1 plays a role in both the onset and maintenance of MLV repression, and its depletion leads to increased expression of the MLV-GFP reporter, suggesting that Smarcad1 is necessary for proper MLV repression in embryonic stem cells.

## Declarations

### Ethics approval and consent to participate

Not applicable

## Consent for publication

Not applicable

## Availability of data and materials

Not applicable

## Authors’ contributions

I.B. and C. S conducted the research. Conceptualization was performed by I.B. and S.S.; methodology was developed by I.B. and C.S.; I.B. drafted the figures; S.S. wrote the main manuscript text and prepared the figures; S.S. was responsible for the acquisition of the financial support, management and coordination of the research. All authors reviewed the manuscript.

## List of abbreviations

MLV: murine leukemia virus
ESC: embryonic stem cells
ERV: endogenous retroviruses
PBSpro/gln: proline / glutamine primer binding site
H3.3: histone 3.3
RT: reverse transcription
ChIP: chromatin immunoprecipitation
qPCR: quantitative polymerase chain reaction

## Acknowledgments

We thank Dr. Jacqueline Mermoud for her help with the design and cloning of the Smarcad1 shRNA, for ChIP advice, and for many insightful and encouraging discussions along the way. We wish to thank Prof. Eran Meshorer for providing the H3.3-HA KH2 cell line. We thank all members of the Schlesinger laboratory for helpful discussions and support.

