## Supplemental Figures and Tables for "The role of Smarcad1 in retroviral repression in mouse embryonic stem cells"

### Supplementary materials:

#### ***Supplementary Figure S1: Smarcad1 is recruited to repress incoming retroviral infection.*** (A) Immunoblotting of shSmarcad1 and shControl ESC extract using Smarcad1 antibody, with anti Gapdh as a loading control. A full blot is shown. (B) Expression levels over time following Smarcad1 depletion were measured by RT‒qPCR (C). The calibration of viral copy numbers was determined by genomic DNA extraction and qPCR analysis using provirus primers on the provirus and normalized to Gapdh and control samples with one integration event. In two different experiments, the copy number was similar between the WT and Smarcad1-depleted cells. n=2, Data are the mean ± s.e.m.

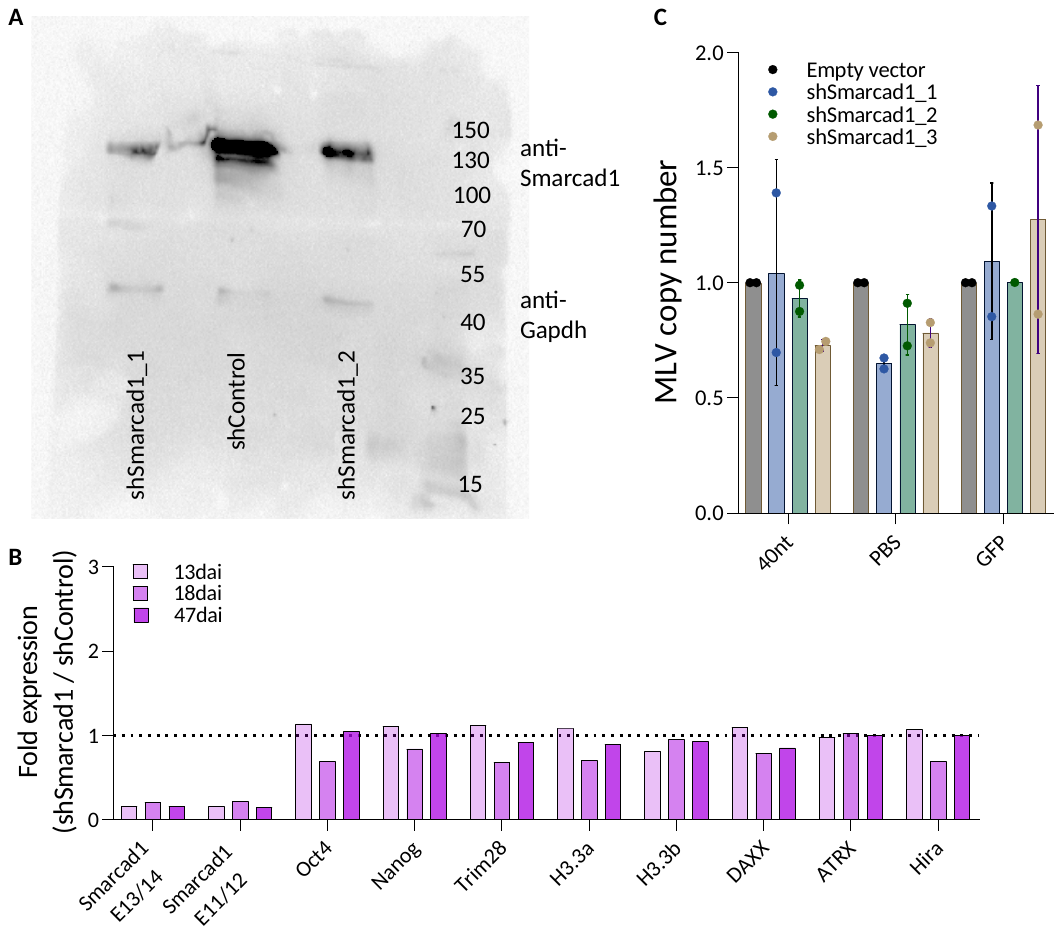

#### ***Supplementary figure S2: Smarcad1 and Trim28 depletion has an additive effect on MLV upregulation.*** *(A) Expression levels of selected ERVs following Smarcad1 and Trim28 depletion were measured by RT‒qPCR. Data are the mean ± s.e.m. (n=2-5) (B) Expression levels of key pluripotency genes and other factors involved in retroviral silencing following Smarcad1 and Trim28 depletion 20 d.a.i. were measured by RT‒qPCR. Data are the mean ± s.e.m. (n=2-5). In all panels, the P value was calculated using a two-tailed unpaired Student's t test, *P<0.05, **P<0.01.*

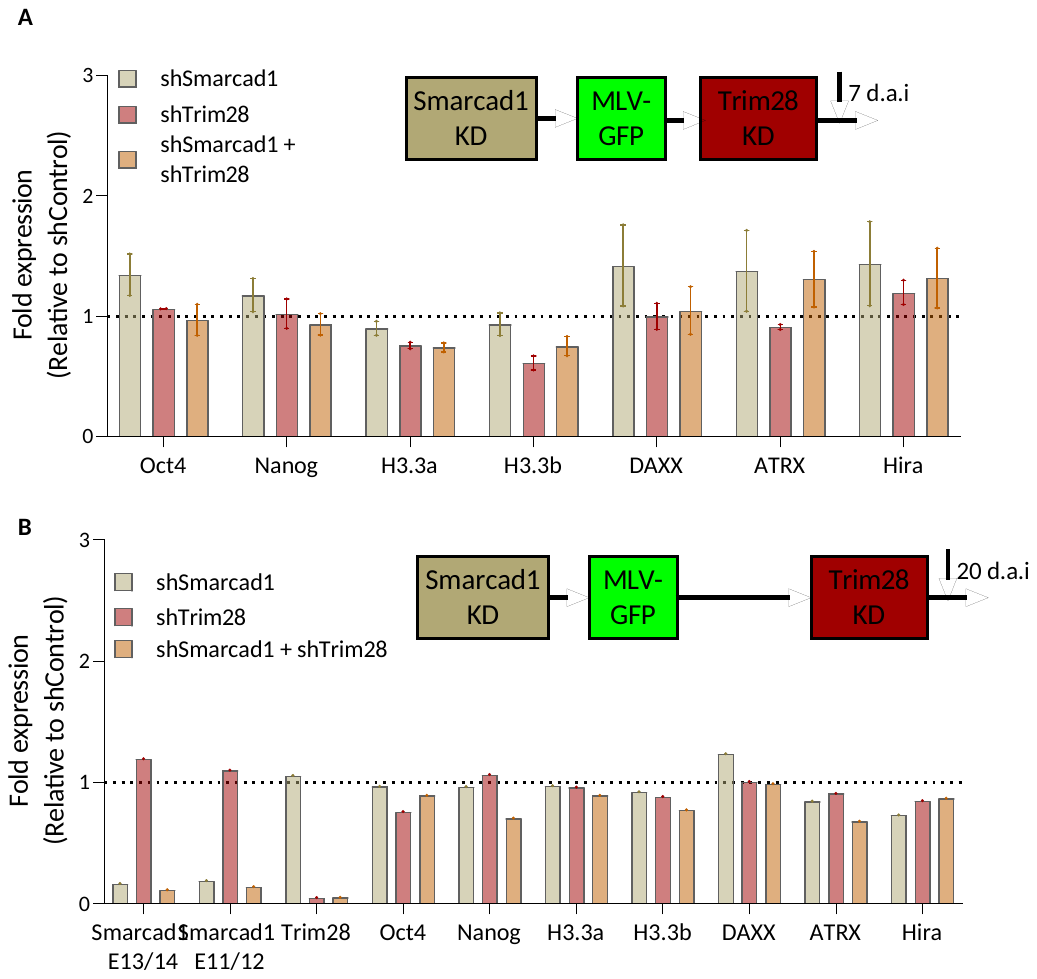

#### Supplementary Table S1: Primers list

| Primer name | Target | 5'-> 3' Sequence | Purpose in this study |
| --- | --- | --- | --- |
| GAPDH_F | GAPDH  (Genomic) | ACCTTTAGCCTTGCCCTTT | ChIP |
| GAPGH_R |  | ACATCACCCCCATCACTCAT |  |
| m-Oct4_pr20 | Oct4  (Genomic) | CACCGGACACCTCACAAAC | ChIP |
| M-Oct4_pr19 |  | TCTCCAGAGGATGGCTGAGT |  |
| L-mChip-Polrmt | Polrmt  (Genomic) | AGACACCTGCTGCCCTATGT | ChIP |
| R-mChip-polrmt |  | GCTCCATCCCAGTGCTTTAC |  |
| Fkbp6_F | FKbp6  (Genomic) | CATGCTCGCTGCGTCTATC | ChIP |
| Fkbp6_R |  | ATCTTGCCGCACAACTGTCT |  |
| HBBb1_F | Hbb  (Genomic) | GCCATAGCCACCCTGTGTAG | ChIP |
| HBBb1_R |  | CTGCCCACTCTGTCCTCTCT |  |
| L-mChip-Peg13 | Peg13  (Genomic) | AGCCTCTGTGCTAGCGTCTC | ChIP |
| R-mChip-Peg13 |  | GGATACCTTCGAGCGTTGAG |  |
| VL30_U3I_LTR _F | VL30 U3I LTR  (Class I ERV) | AGATGTATTGCCAAACACAGG | ChIP/qPCR |
| VL30_U3I_LTR_R |  | AGGGGGAATGGGGAGGGAA |  |
| RLTR1b_F | RLTR1b  (Class I ERV) | TCCTTCCCTTTGCCCTATTT | ChIP/qPCR |
| RLTR1b_R |  | GGCTGGAACTGGTGAGATGT |  |
| RLTR45_F | RLTR45  (Class II ERV | TGCTTTTCCGACATGGTAAT | qPCR |
| RLTR45_R |  | AGTAACCCTGACCTGCTCCT |  |
| ETnERV2_F | EtnERV2  (Class II ERV) | ACAAATTCAGTATGGGCATC | ChIP/qPCR |
| ETnERV2_R |  | GGGTACTGTTAAGACCCACA |  |
| IAP-F | IAP  (Class II ERV) | AAGCAGCAATCACCCACTTTGG | ChIP/qPCR |
| IAP-R |  | CAATCATTAGATGTGGCTGCCAAG |  |
| IAP1-MM_I-F | IAP1 MM I  (Class II ERV) | CGTTGCCTTGAGTTTCAACA | ChIP/qPCR |
| IAP1-MM_I-R |  | GTCACATCCATCTGCCACAC |  |
| IAP5-Mm_I-F | IAP5 MM I  (Class II ERV) | GTGCGGTCAGTCCTACGTAA | ChIP/qPCR |
| IAP5-Mm_I-R |  | CCCGGAGCAGAAGTGAAAGT |  |
| IAPEY3_i-F | IAPEY3i  (Class II ERV) | TGGTGTGGAAGATGCTGACC | ChIP/qPCR |
| IAPEY3_i-R |  | CCCATCTCTGTTCGCCCATT |  |
| IAPEY4_i-F | IAPEY4i  (Class II ERV) | TCATCGAGCCAGTGAGTGTC | ChIP/qPCR |
| IAPEY4_i-R |  | TTCTGCCTGAGGTCCAGACT |  |
| IAPEY5_I-F | IAPEY5i  (Class II ERV) | GACAATGGGCCAGCTTATGT | ChIP/qPCR |
| IAPEY5_I-R |  | GCCTTGTCCTTGTGGATTGT |  |
| IAPEz_F_ChIP | IAPEz  (Class II ERV) | GCTCCTGAAGATGTAAGCAATAAAG | ChIP/qPCR |
| IAPEz_R_ChIP |  | CTTCCTTGCGCCAGTCCCGAG |  |
| IAPEz-int_F | IAPEz int  (Class II ERV) | CCATCTTGTAACGGCGAATGT | ChIP |
| IAPEz-int_R |  | AGCAATAGGTATGGTGGTA |  |
| IAPEz-gag_F | IAPEz gag  (Class II ERV) | CACGCTCCGGTAGAATACTTACAAAT | ChIP |
| IAPEz-gag_R |  | CCTGTCTAACTGCACCAAGGTAAAAT |  |
| IAPLTR1a_I_MM-F | IAPLTR1a  (Class II ERV) | CAGTGCGCAGACTCATTCAT | ChIP |
| IAPLTR1a_I_MM-R |  | CTCGGCTCCTTCAAAGACTG |  |
| IAPLTR3_I-F | IAPLTR3i  (Class II ERV) | GGCAAACCTCAGAAGGACAG | ChIP |
| IAPLTR3_I-R |  | CACATCTCTGCCCCATAGGT |  |
| IAPLTR4_I-F | IAPLTR4i  (Class II ERV) | CAGCTGAAAGGCACAGACAA | qPCR |
| IAPLTR4_I-F |  | AAATAGGATCCGGGCCATAC |  |
| IAP gag F | IAP gag  (Class II ERV) | AATCTCAGAACCGCTCCATGA | ChIP/qPCR |
| IAP gag R |  | TTTCTTAAAATGCCCAGGCTTT |  |
| IAP pol F | IAP pol  (Class II ERV) | CTTGCCCTTAAAGGTCTAAAAGCA | ChIP |
| IAP pol R |  | GCGGTATAAGGTACAATTAAAAGATA |  |
| MMERVK-10C_F | MERVK10C  (Class II ERV) | CAAATAGCCCTACCATATGTCAG | ChIP |
| MMERVK R |  | GTATACTTTCTTCTTCAGGTCCAC |  |
| MERVL-F | MERVL  (Class III ERV) | ATCTCCTGGCACCTGGTATG | ChIP/qPCR |
| MERVL-R |  | AGAAGAAGGCATTTGCCAGA |  |
| Line1_1 ChIP_F | Line1  (Non-LTR) | GATTACCAGATGGCGAAAGG | ChIP/qPCR |
| Line1_ChIP_R |  | AGTGCTGCGTTCTGATGATG |  |
| PBS_F | MLV PBS  (Provirus) | TGGCCAGCAACTTATCTGTG | ChIP |
| PBS_R |  | CAGGCGCATAAAATCAGTCA |  |
| GFP2-F | MLV GFP  (Provirus) | AGCTGAAGGGCATCGACTT | ChIP |
| GFP2-R |  | GATGCCGTTCTTCTGCTTGT |  |
| MLV +4-F | MLV TSS  (Provirus) | CAGTCCTCCGATTGACTGAG | ChIP |
| MLV +91-R |  | GGAACAGCGAGACCACAAGT |  |
| 40nt_L | MLV 40nt  (Provirus) | CCTCCACACACCATCATCCA | ChIP |
| 40nt2_R |  | CCTGTTTGGCCCATATTCAG |  |
| UBC E1 L | Ubiquitin C  (UbC) | CAGCCGTATATCTTCCCAGACT | qPCR |
| UBC E2 R |  | CTCAGAGGGATGCCAGTAATCTA |  |
| TRIM28 E4 L | Trim28  E4/5 | CCTCGTGCTGTTCTGTGAGA | qPCR |
| TRIM28 E4/5 R |  | TGGTACTGATGGTCCTTGTGA |  |
| SMARCAD1 E11/12 F | Smarcad1  E11/12 | AGACGAAATGGGCCTAGGAA | qPCR |
| SMARCAD1 E12 R |  | GTGGAGGCTGGAACAACAAT |  |
| SMARCAD1 E13 | Smarcad1  E13/14 | TCCACCATAGATAACTGGTTAAGAGA | qPCR |
| SMARCAD1 E13/14 |  | TGAGAACCATAGTAACAGAGCACAT |  |
| H3.3A E1/2 F | H3.3a  E2/3 | CAGCGCCGCCTCTCGCTTG | qPCR |
| H3.3A E1/2 R |  | CAGTCTGCTTTGTACGAGCCATGGTA |  |
| H3f3B-E2-E3-F | H3.3b | CCAAGGCGGCTCGGAAAAGC | qPCR |
| H3f3B-E2-E3-R |  | GGTAACGACGGATCTCTCTCAGA |  |
| HIRA E7 F | HIRA  E7/9 | AACTCTGAGAGGTCATTCTG | qPCR |
| HIRA E9 R |  | CACAACAGTCACAGCTTTCC |  |
| DAXX_E3/4 | DAXX | CTATAGGCCAGGCGTTGACC | qPCR |
| DAXX_E4 |  | GCCAAGGTTCGATTTTCCCG |  |
| shRNA Forward Primer | pLKO.1 multiple cloning site | ATCGTTTCAGACCCACCTCC | Cloning |
| shRNA Reverse Primer |  | CAGGGCTGCCTTGGAAAAG |  |

Supplementary Table S2: Antibodies used in this study.

| Species in which antibody was produced | Target | Clonality | Stock concentration | Dilution of stock for WB | Source, cat number | Use in lab |
| --- | --- | --- | --- | --- | --- | --- |
| Mouse | Trim28 | Monoclonal | 1mg/ml | N/A | Abcam, ab22553 | ChIP |
| Rabbit | HA tag | Polyclonal | 1mg/ml | 1:1000 | Abcam, ab9110 | ChIP & WB |
| Rabbit | Smarcad1 | Polyclonal | N/A | 1:1000 | Sigma, HPA016737 | ChIP & WB |
| Rabbit | GAPDH | Polyclonal | 200 µg/ml | 1:1000 | Santa Cruz, sc-25778 | WB |
